# Genetic differentiation is constrained to chromosomal inversions and putative centromeres in locally adapted populations with higher gene flow

**DOI:** 10.1101/2024.10.20.619329

**Authors:** Maria Akopyan, Anna Tigano, Arne Jacobs, Aryn P. Wilder, Nina O. Therkildsen

**Author notes:** **Correspondence:** Maria Akopyan, **Email:**. Current affiliation: Department of Quantitative and Computational Biology, University of Southern California, Los Angeles, California, USA.

## Abstract

The impact of genome structure on adaptation is a growing focus in evolutionary biology, revealing an important role for structural variation and recombination landscapes in shaping genetic diversity across genomes and among populations. This is particularly relevant when local adaptation occurs despite gene flow, where clustering of differentiated loci can maintain locally adapted variants by reducing recombination between them. However, the limited genomic resources for non-model species, including reference genomes and recombination maps, has constrained our understanding of these patterns. In this study, we leverage the Atlantic silverside—a non-model fish with extensive local adaptation across a steep latitudinal gradient—as an ideal system to explore how genome structure influences adaptation under varying levels of gene flow, using a newly available reference genome and multiple recombination maps. Analyzing 168 genomes from four populations, we found a continuum of genome-wide differentiation increasing from south to north, reflecting higher connectivity among southern populations and reduced gene flow at northern latitudes. With increasing gene flow, the number and clustering of F_ST_ outlier loci also increased, with differentiated loci tightly clustered in large haploblocks harboring inversions and smaller peaks overlapping putative centromeres. Notably, sequence divergence was only evident in inversions, supporting their role in adaptive divergence with gene flow, whereas centromeres appeared differentiated because of low recombination and reduced diversity, with no indication of elevated sequence divergence. Our results support the hypothesis that clustered genomic architectures evolve with high gene flow and enhance our understanding of how inversions and centromeres are linked to different evolutionary processes.

**Significance Statement:** How populations preserve favorable combinations of genes adapted to their local environment despite reproducing with populations adapted to different conditions is a longstanding question in evolutionary biology. By analyzing the genomes of 168 Atlantic silverside fish from four populations, we found that when populations adapted to different environments frequently interbreed, genetic differences concentrate in specific parts of the genome, particularly in chromosomal inversions—where segments of DNA are flipped. These inversions help preserve locally adapted gene combinations, enabling populations to maintain differences essential for survival in their habitats. This research enhances our understanding of genomic adaptation, a fundamental evolutionary question with increasing relevance as environmental changes pose new challenges globally.

## Introduction

Understanding the complex interplay between natural selection and gene flow is crucial for discerning how adaptations evolve and are maintained. Populations can adapt to local environmental conditions in response to selection, but this process may be hindered if populations under divergent selection are still connected by gene flow (1, 2). Because gene flow can introduce maladaptive alleles and homogenize populations, it could swamp adaptation, depending on the strength of selection and the degree of connectivity (3–5). Developments in genomic techniques and evolutionary simulations in recent decades have led to a growing appreciation for the role of genomic architecture underlying adaptive traits – including structural variation, chromosomal organization, and recombination landscapes – in local adaptation with gene flow (6, 7). While the role of structural variation in adaptation is increasingly recognized (8–10), the broader influence of genome structure on the distribution of genetic variation across genomes and among populations, and its interaction with various evolutionary processes, is still poorly understood.

The recombination landscape, i.e. the variation in recombination rates across the genome, emerges as a key player in the dynamic interplay between selection and gene flow (11–13). Regions of relatively low recombination may offer protection from the deleterious effects of maladaptive gene flow and allow populations to maintain clusters of locally adapted alleles (14–16). For instance, recombination modifiers such as chromosomal inversions – mutations that change the orientation of a DNA segment within a chromosome – are increasingly implicated in studies of adaptation and divergence (reviewed in 17). When individuals inherit both arrangements of a chromosomal inversion, they often produce non-viable gametes if recombination occurs between these inverted regions of the genome (18, 19). Consequently, recombination in these regions is heavily reduced between alternate arrangements at the population level, often resulting in strongly linked alleles within an inversion that can act similarly to a single large-effect locus. This in turn allows populations, particularly in the face of maladaptive gene flow, to maintain sets of locally adapted alleles at high frequency (20).

Areas of suppressed recombination that do not coincide with chromosomal inversions may also play a role in adaptation with gene flow, but have received far less attention. Centromeres, which are essential for chromosome segregation during cell division, often exhibit reduced recombination rates, impacting the distribution of genetic diversity along chromosomes. For instance, sequence divergence between cryptic species of alpine bumblebees is elevated in regions of low recombination and near centromeres via genetic hitchhiking, with accentuated divergence in these regions in the presence of gene flow (21). However, few studies have examined the role of centromeres in adaptive divergence, and in most cases, the positions of centromeres are unknown, in part due to the difficulty of sequencing and assembling regions of highly repetitive DNA. In addition, whereas chromosomal inversions only suppress recombination in a heterozygous state (18), recombination in centromeres is consistently low (Fig 1A). As a result, inversions may be more likely to facilitate adaptive divergence in the face of gene flow, compared to centromeric loci where consistently low recombination decreases the likelihood of incorporating adaptive variants and purging deleterious ones. Despite fundamental differences in how centromeres and inversions modify the recombination landscape, studies distinguishing their impacts on the genomic patterns of adaptive divergence are lacking.

**Figure 1.**
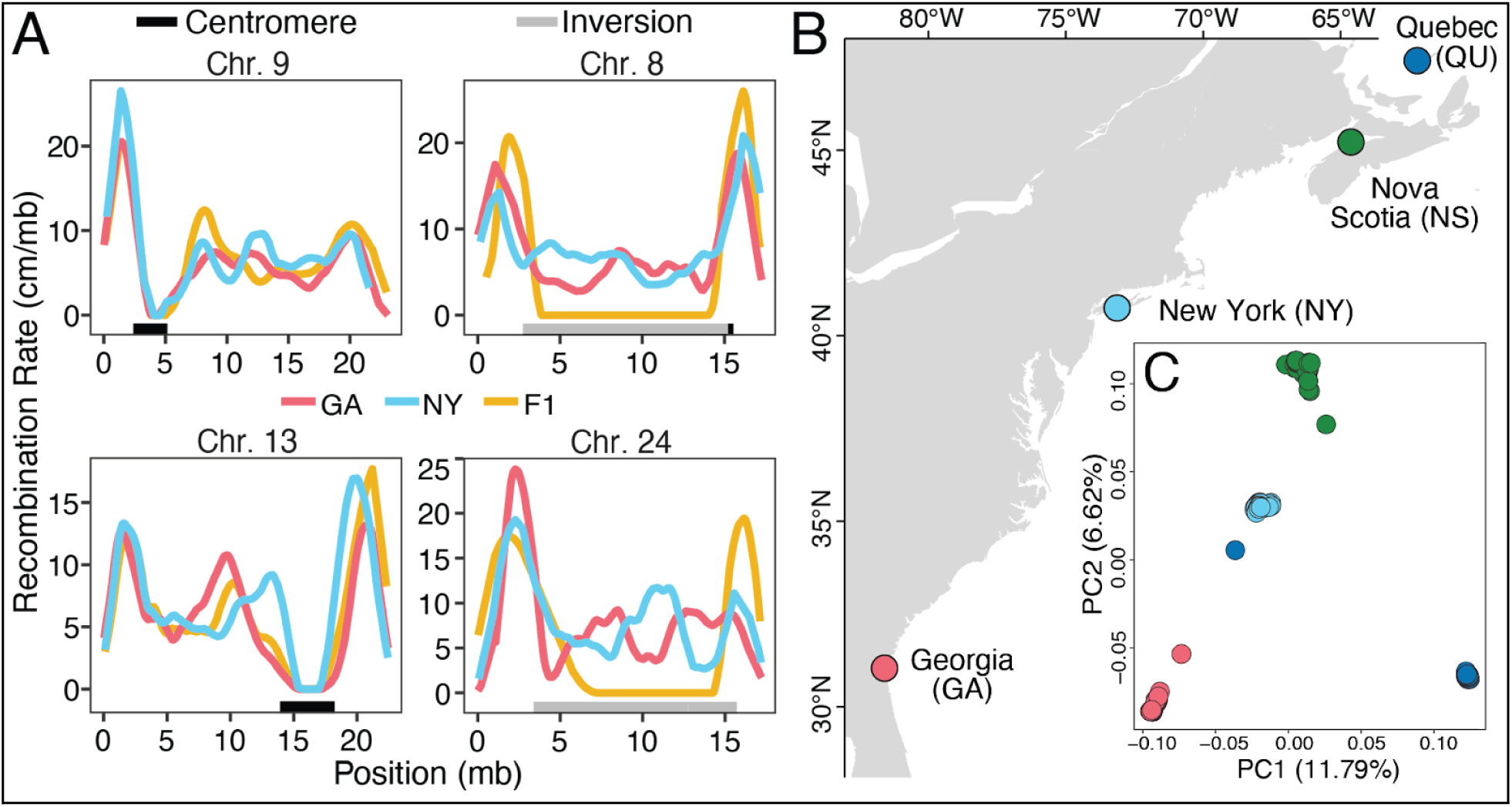
A) Recombination rates within and between GA (red) and NY (blue) populations and their interpopulation hybrids (gold) across representative chromosomes from Akopyan et al. (61). Horizontal bars below plots indicate inversions (gray) and putative centromeres (black). Recombination rate drops to near zero in all three maps in the putative centromere regions, while the recombination rate in inversion regions stays at intermediate levels within the pure-population maps (GA and NY) and only drops to near-zero in the map for the inter-population hybrid. B) Sampling localities of silverside genomes obtained from Wilder et al. (59). Collection sites include Jekyll Island, Georgia (GA), Patchogue, New York (NY), Minas Basin, Nova Scotia (NS), and Magdalen Island, Quebec (QU). C) Principal components 1 and 2 of genome-wide variation among individuals, colored by sampling locations on the map in panel B.

Variation in recombination rates may also be associated with various features of the genome, for instance, at broad scales with respect to genome and chromosome size (22, 23), and at finer scales with respect to nucleotide, repeat, and genic content (24). Within chromosomes, recombination events are likely influenced by the tendency for double-strand breaks to occur near the telomeres where chromatin is open and accessible. In contrast, as mentioned above, recombination tends to be inhibited within and around the centromeres, due to high heterochromatin density (25, 26). Additionally, a strong correlation between GC content and recombination rate has been shown in fish (27), birds (28) and mammals (29, 30), a pattern resulting from GC-biased gene conversion, by which fixation of alleles containing G and C bases is accelerated relative to A and T bases due to their stronger molecular bonds (31). Furthermore, recombination occurs more frequently in gene-dense regions in mice (32) and plants (33, 34), and in the vicinity of, but not within, coding and regulatory regions in insects (35–37), and humans (38). Despite recombination landscapes having been characterized in multiple taxa, detailed recombination maps are still primarily available for model organisms and domesticated species, making it difficult to assess the extent to which regional variation in recombination rate is associated with genomic features, the efficiency of selection, and the levels of genetic diversity in natural populations across species.

Commonly used population genetics statistics, including differentiation (F_ST_) and nucleotide diversity (π), are heavily influenced by the variation in recombination rate across the genome (39–42). These relationships make it hard to distinguish whether blocks of elevated differentiation identified between populations are a result of divergence with gene flow, and in fact underpin local adaptation, or result from other selective pressures interacting with the landscape of recombination. Because regions with low recombination rates experience stronger effects of linked selection, resulting in selection impacting a larger contiguous region of the genome, both purifying selection (i.e. background selection) and positive selection in isolation can emulate the peaks of differentiation caused by divergence with gene flow. However, sequence divergence (d_XY_), an absolute measure of differentiation that is not dependent on within-population variation, is expected to differ in these alternate scenarios. Under the divergence with gene flow model, the regions that contribute to divergence between populations are expected to show elevated d_XY_ compared to background levels of absolute differentiation, whereas in isolation, selection should reduce or have no effect on d_XY_ compared to background levels (40). While some studies, primarily focused on birds, have examined the signatures resulting from these different models of selection (43–47), empirical studies explicitly investigating differences among recombination-modifying regions of the genome such as inversions and centromeres are still lacking, and most of what we know about the role of recombination in the evolution of clustered architectures of adaptation and divergence is based on simulations (16, 48–50). The need for high-quality genomic resources, including chromosome-level reference genomes and recombination maps, combined with the need for a so-called natural laboratory, i.e. an example of adaptive divergence in multiple traits that have evolved in populations with and without gene flow, have likely contributed to the delayed development of this research area.

The Atlantic silverside, *Menidia menidia*, a small fish distributed across the steep latitudinal climate gradient of the North American Atlantic coast, is an excellent system to examine the role of recombination modifiers in adaptive divergence with gene flow. Extensive research on this species has demonstrated a remarkable degree of local adaptation across its clinal range in multiple complex traits (51), likely underpinned by many contributing genes. Atlantic silversides exhibit countergradient variation in growth rate (52), with populations at northern latitudes growing faster to compensate for shorter growing seasons, and southern populations growing slower due to tradeoffs with predator avoidance (53–55). Silverside populations also exhibit clinal genetically based variation in other complex traits, including vertebral number, swimming performance, temperature-dependent sex determination, lipid storage, spawning temperature and duration, and offspring size at hatch (reviewed in 56). Interestingly, adaptive differences are maintained within populations despite evidence of high dispersal abilities (57). Additionally, genetic data supports the presence of three regional population groups, with higher levels of connectivity among southern populations and reduced connectivity at northern latitudes (58), providing an opportunity to study patterns of adaptive divergence across varying levels of gene flow.

Recent exome-based population genomics work in Atlantic silversides suggested that geographic differentiation was primarily concentrated in large blocks on multiple chromosomes, but also revealed more scattered signatures of differentiation across the genome (59). However, the lack of a species reference genome limited inference of exact genomic positions of outlier regions and of the association between elevated differentiation and the underlying features of the genome. Leveraging a high-quality chromosome-level reference genome for the Atlantic silverside (60) and linkage maps (61), we revisit a large low-coverage whole-genome resequencing data set (59) to characterize patterns of population genetic diversity and differentiation in light of the underlying recombination landscape and assess potential associations between the recombination landscape and nucleotide composition and genomic features. We also calculate an absolute measure of sequence divergence between populations to distinguish the genomic regions with evidence of reduced gene flow between populations, as such regions likely contain loci that contribute to adaptive divergence with gene flow (40). We test the hypothesis that higher gene flow will favor concentrated architectures of differentiation, predicting that the clustering of differentiated alleles in low recombining regions increases with increasing levels of gene flow between populations. Further, we examine whether patterns of diversity, differentiation, and divergence differ between conditionally low-recombining inversions and consistently low-recombining centromeres. Our study is one of the first to empirically test these fundamental predictions about the relative roles of centromeres and inversions in adaptation with gene flow.

## Results

### Levels of background differentiation (F_ST_) decrease from north to south

We analyzed whole genome sequence data from 42 fish from each of four populations (168 individuals total) along the North American Atlantic coast to assess population structure and differentiation (Fig 1B). A total of 413,215,831 sites in the reference genome (including variant and invariant sites) passed our quality filters, representing 89% of the genome that is assembled into chromosomes, and we identified 20,421,651 SNPs across all 168 individuals with an average depth per individual of 1.4×. The four populations formed distinct clusters along principal components of genome-wide variation, mirroring the geographic distribution of populations, with the exception of one individual from the north clustering with southern populations (Fig 1C). The northernmost population (QU) showed the most separation in the PCA relative to the rest of the populations, reflecting the history of independent colonization of this region from the southern coastline and low ongoing gene flow (58). Pairwise comparisons of neighboring populations revealed the lowest levels of average genome-wide differentiation between GA and NY populations (mean F_ST_=0.023, median F_ST_=0.01), increasing two-fold between NY and NS (mean F_ST_=0.040, median F_ST_=0.023), and three-fold between NS and QU (mean F_ST_=0.068, median F_ST_=0.031). Pairwise allele frequency difference (AFD), a metric similar to F_ST_ that is more linear and sensitive to weak population differentiation (62), also suggests overall high gene flow in the south and a continuum of connectivity among populations: median AFD increases from 0.043 to 0.066 to 0.084 when comparing neighboring populations from the south to the north.

### Populations with higher gene flow exhibit clustering of F_ST_ outliers

We examined F_ST_ outlier regions across chromosomes to investigate differentiation patterns between neighboring populations (Fig 2). Between the two southernmost populations, clusters of F_ST_ peaks are prominent, in contrast to low levels of genome-wide differentiation (Fig 2C). These results corroborate previous exome-based work which revealed that differentiation was primarily concentrated in blocks (Fig S1, 59). We found this pattern to be even more pronounced with low-coverage whole-genome data mapped to the species-specific reference genome. Many of the peaks that appeared to be scattered across the reference transcriptome anchored to the medaka genome now fit into larger blocks when mapped to the silverside genome (Fig 2, 59).

**Figure 2.**
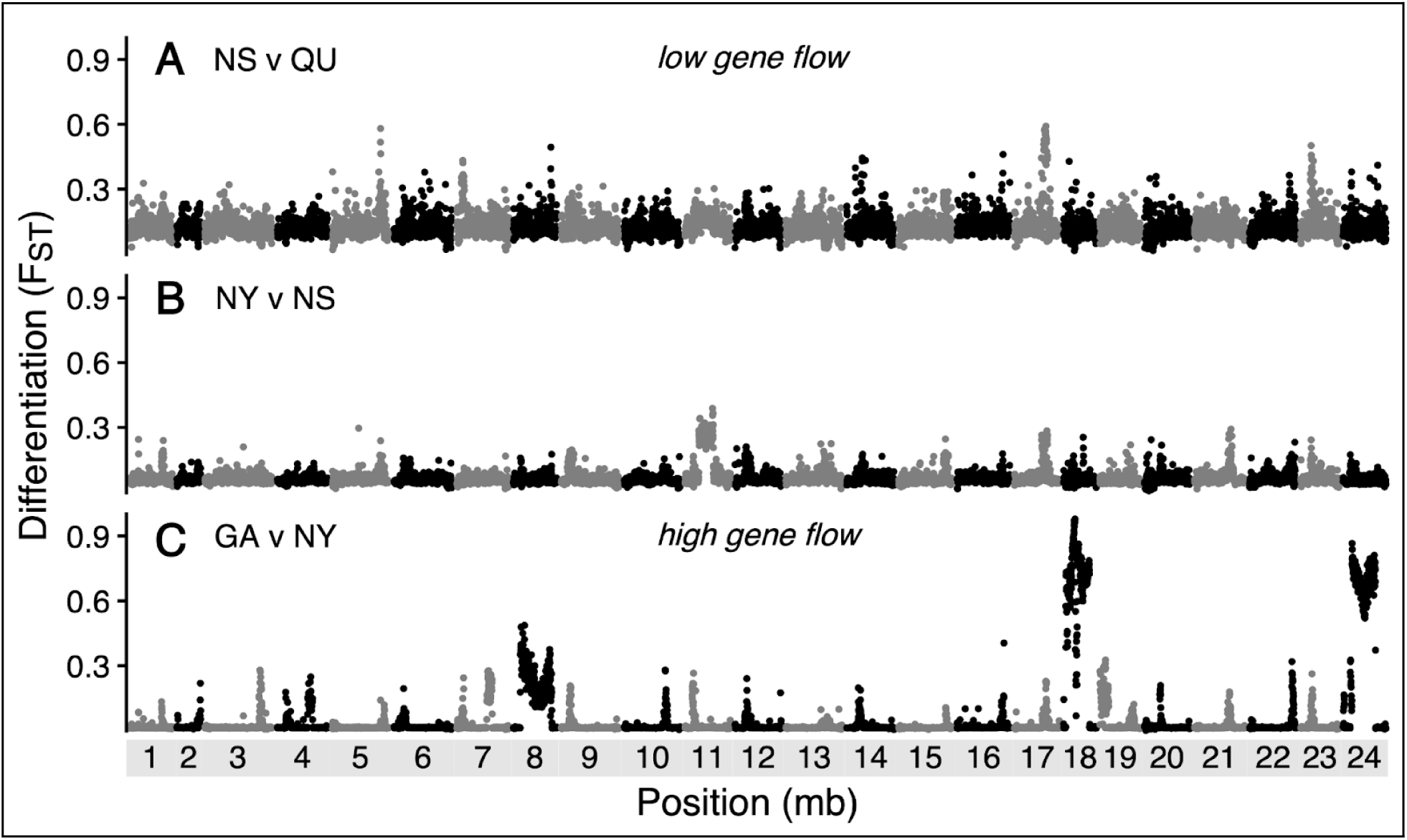
Genome-wide distribution of F_ST_, averaged in 50 kb windows, across the 24 chromosomes of the Atlantic silverside genome, for pairwise comparisons of neighboring populations as indicated in Fig 1B. Comparison between A) Nova Scotia (NS) and Quebec (QU), where gene flow is limited, B) between New York (NY) and NS, and between C) Georgia (GA) and NY, where gene flow is high, demonstrating clustering of F_ST_ peaks among low levels of genome-wide differentiation

In the southern population comparisons where we see lower background differentiation, likely because of higher gene flow, we identified more outlier regions (50 kb windows) with elevated differentiation on fewer chromosomes compared to the northernmost populations. Comparing populations from QU and NS (northern), we identified 150 outlier F_ST_ regions distributed across 21 chromosomes. Between NS and NY (mid-latitude), we identified 215 outlier F_ST_ regions distributed across 20 chromosomes, and between NY and GA (southern), we identified 360 outlier F_ST_ regions distributed across 3 chromosomes: 8, 18, and 24. To assess whether outliers were non-randomly clustered in the genome, we performed a permutation test by redistributing outliers across chromosomes (weighted by size) 10,000 times and compared the observed variance to a null distribution. In all three comparisons, F_ST_ outlier windows were unevenly distributed across chromosomes, with the observed concentrations deviating significantly from expected random distributions (p<0.0001), indicating a strong clustering of outliers on certain chromosomes, most notably in the NY vs. GA comparison. The observed variances were approximately 4, 33, and 203 times larger than simulated means for the comparisons between QU and NS, NS and NY, and NY and GA, respectively, highlighting that as gene flow increases, F_ST_ outliers become intensely clustered on specific chromosomes, with the most extreme clustering seen in the NY vs. GA comparison where gene flow is highest (Fig 2C).

### Massive chromosomal inversions harbor haploblocks of differentiation

We identified the same four major haploblocks of highly elevated differentiation described previously (59): a region on chromosome 11 in the pairwise comparison of NY and NS (Fig 2B), and regions on chromosomes 8, 18, and 24 in the comparison of NY and GA (Fig 2C). The largest haploblock with elevated differentiation was observed on chromosome 8 and covered 12.5 Mb, which represents 72% of the assembled chromosome length. The haploblocks on chromosomes 18 and 24 spanned 9.6 Mb and 9.4 Mb, representing 73% and 54% of the respective assembled chromosomes. The haploblock between NY and NS on chromosome 11 was 8.6 Mb long and spanned 47% of the chromosome. Between NY and GA, a narrow F_ST_ peak was observed on chromosome 11 coinciding with one of the endpoints of the haploblock present in the north (Fig 2). The number of nearly fixed variants (F_ST_>0.95) revealed a striking contrast in magnitude across population comparisons, with a single variant identified between NS and QU and between NS and NY, located on chromosomes 5 and 11, respectively. In contrast, GA vs. NY shows a total of 55,806 nearly fixed variants, with dramatic peaks on chromosomes 18 (25,454 variants) and 24 (30,009 variants), as well as a notable cluster on chromosome 8 (336 variants), underscoring significant differentiation in these regions.

Average F_ST_ per 50 kb window significantly decreased with increasing recombination rate (Fig S2), especially recombination rate between lab-reared interpopulation crosses (τ=-0.21, p<0.0001), and low recombination between hybrids was a hallmark of the highly differentiated peaks and haploblocks between GA and NY (Fig 3). The four large haploblocks on chromosomes 8, 11, 18, and 24 coincide with known massive chromosomal inversions where linkage mapping has shown that recombination is suppressed between alternate arrangements (61). Peaks of differentiation on chromosomes 4, 7, and 19 between southern populations also overlap known segregating inversions (Fig 3). Further, PCA analysis revealed a clustering pattern typical of inversions, with three main clusters corresponding to the two inversion homozygotes and the heterozygotes, consistent with the suppression of recombination within these regions (Fig S3).

**Figure 3.**
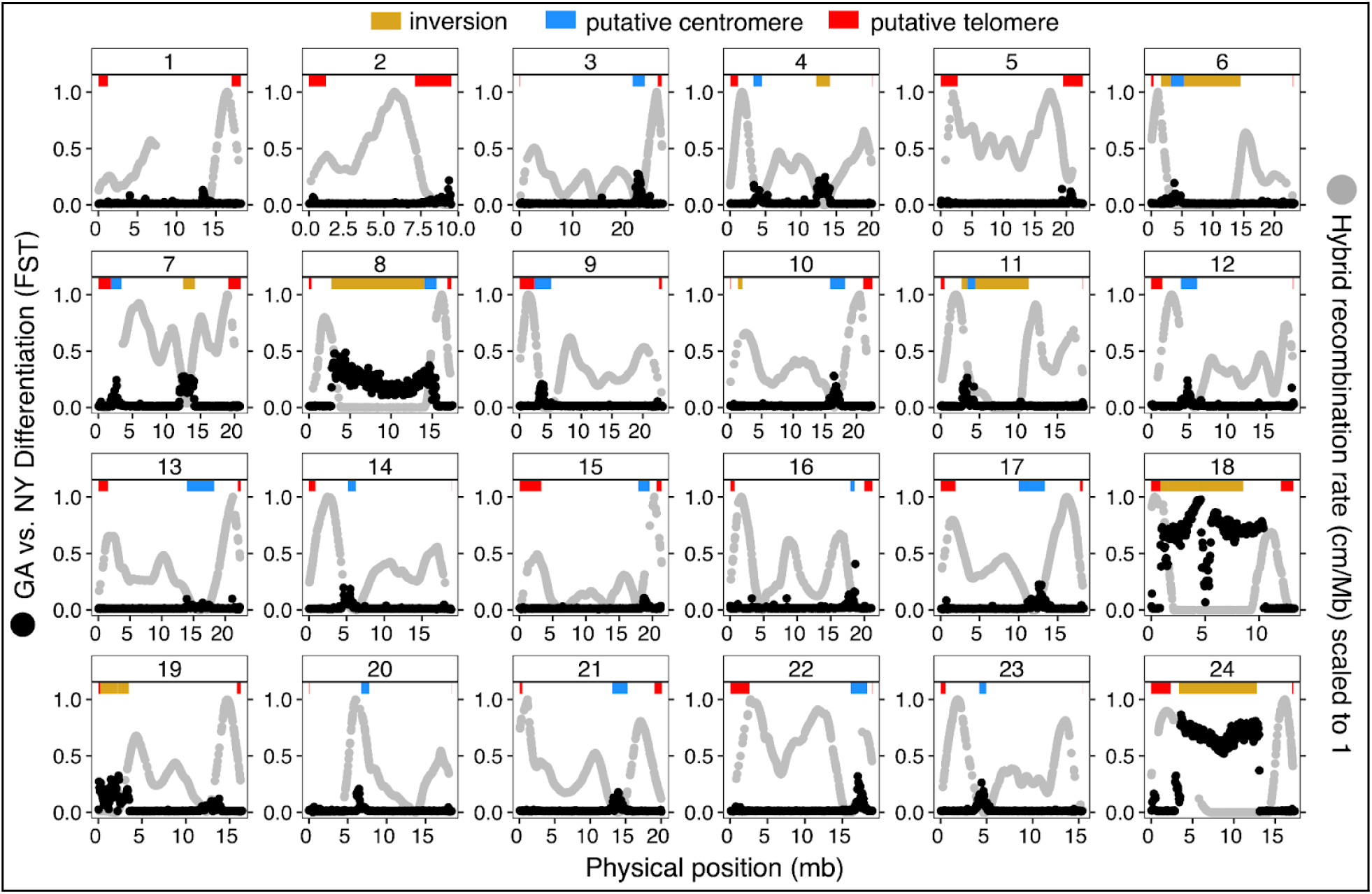
Inversions and putative centromeres coincide with patterns of genomic differentiation and recombination suppression between populations. Black points show F_ST_ in 50 kb windows between the GA and NY populations (left y-axis), and gray points show the recombination rate in their inter-population hybrids (right y-axis) for each chromosome. Putative centromere and telomere positions are indicated by blue and red horizontal bars, respectively, and inversions are indicated by gold bars.

### Localized F_ST_ peaks align with putative centromeres

The pairwise comparison of the two southernmost populations NY and GA showed that most chromosomes have one, sometimes two, prominent peaks in differentiation (Fig 2C). In contrast to the abrupt jump in differentiation observed at the ends of the inversion-associated haploblocks, these peaks had a more mountain-like shape, characterized by a rapid but continuous increase of differentiation towards their centers. They were evident on all chromosomes except those with large haploblocks (chromosomes 8, 18, and 24). In contrast, chromosomes 4, 7, and 19 each had two peaks. Pairwise comparisons involving the northern populations also revealed similar peaks of differentiation, but the pattern was less prominent against higher levels of background differentiation and less consistent across chromosomes. The peaks that do appear, however, were in the same chromosomal positions across population comparisons (Fig 2). For instance, the peaks on chromosomes 17 and 23 appear in all comparisons, the peak on chromosome 21 appears in the pairwise comparisons including GA, NY, and NS, and one of the peaks on chromosome 7 appears only in comparisons involving the southernmost or northernmost neighboring populations.

Strikingly, the majority of the narrow peaks coincided with putative centromeres (Fig 3). We identified putative centromeric regions for 18 out of the 24 chromosomes, and in all 18 centromeric regions, a discernible peak in differentiation was seen between the southern populations where genome-wide differentiation was lowest (Fig 3). With increasing levels of background differentiation between NY and NS and even more so between NS and QU, only 12 and 7 of the 18 centromeric regions had an F_ST_ peak that was detectable by eye, respectively (Fig 2). In all pairwise comparisons of neighboring populations, however, F_ST_ was significantly higher at putative centromeres compared to the rest of the genome (t=20.54, p<0.0001; Fig 4A). Median F_ST_ in centromeric regions was 2.5 times higher between GA and NY, 1.6 times higher between NY and NS, and 1.2 times higher between NS and QU compared to the median background differentiation in each respective pairwise comparison (Wilcoxon test, p<0.001). Telomeres, on the other hand, had significantly lower F_ST_ compared to the rest of the genome between NS and QU (t=9.56, P<0.0001) and between NY and NS (t=7.85, p<0.0001), while between GA and NY, telomeres showed significantly higher F_ST_ (t=-2.68, p=0.007); however, these differences were relatively small and not clearly distinguishable when plotted (Fig 4A). For both centromeres and telomeres, F_ST_ varied not only among population comparisons, but also among chromosomes, with notably more variation across chromosomes in centromeric regions compared to telomeric regions (Fig S3). In addition, PCA of centromeric regions showed similar clustering to inversions but with a much broader spread of individuals, reflecting higher variation and lower differentiation in centromeres compared to inversions (Fig S3).

**Figure 4.**
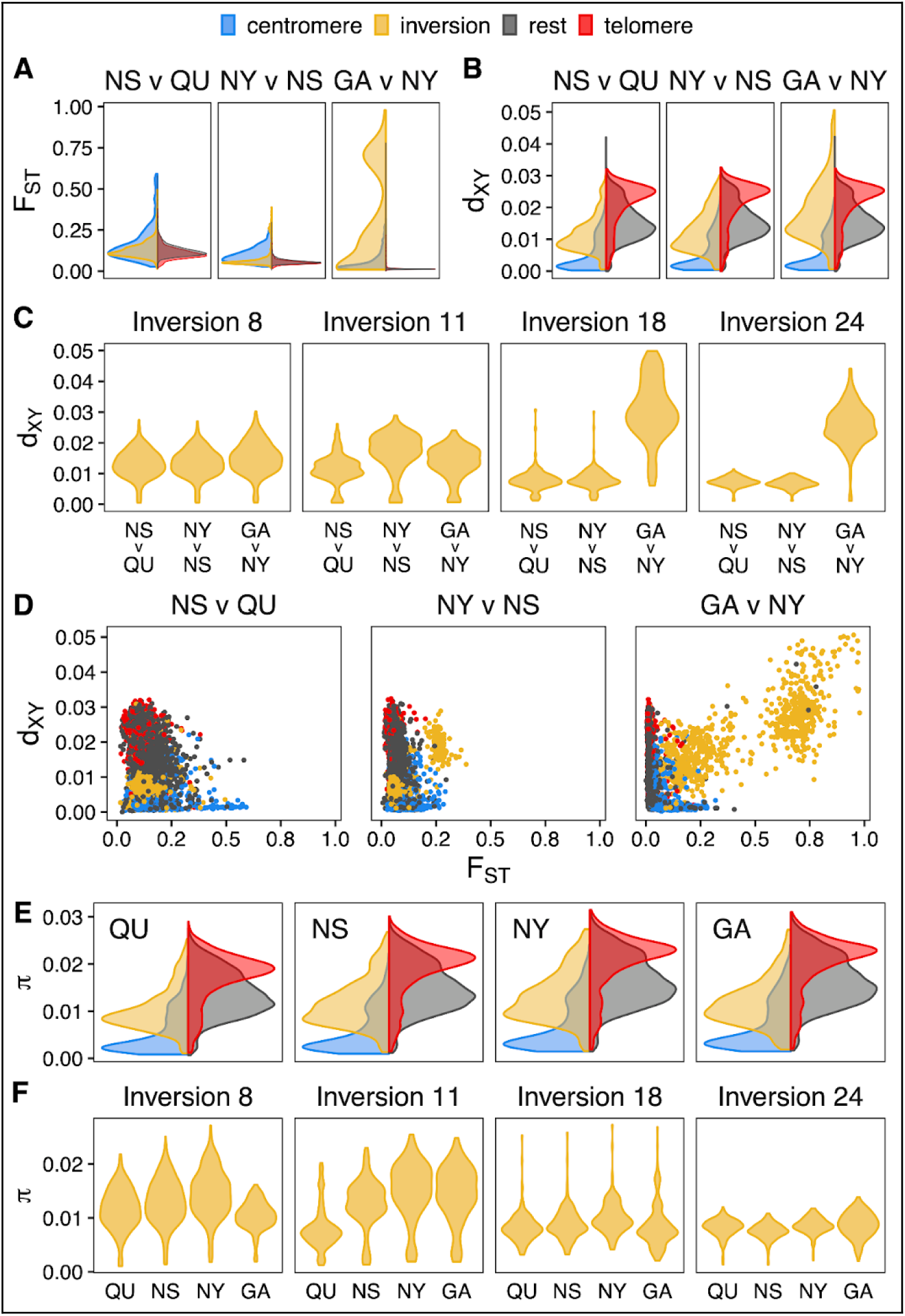
Comparison of diversity (π) and differentiation (F_ST_ and D_XY_) across genomic regions between neighboring populations. Half violin plots display average values in 50 kb windows for centromeres (blue), inversions (yellow), telomeres (red), and the rest of the genome (dark gray) for F_ST_ (A) and d_XY_ (B). In panel (A), the F_ST_ averages for telomeres and the rest of the genome are similar, leading to overlapping half violins that are visually indistinguishable. Relationships between F_ST_ and D_XY_ across 50 kb windows are shown by genomic region (C). π averaged in 50 kb windows across four populations is compared across genomic regions (D) and within each major inversion linked to haploblocks of elevated F_ST_ (E). Panel (F) presents d_XY_ distributions within each major inversion for neighboring population pairs.

### Sequence divergence (d_XY_) is elevated in large chromosomal inversions

Patterns of genome-wide d_XY_ showed dramatic oscillations within chromosomes, dipping low in putative centromeres and peaking at putative telomeres for the majority of chromosomes, i.e., the opposite of the F_ST_ pattern described above (Fig S4). In all pairwise comparisons of neighboring populations, d_XY_ was significantly higher at telomeres (t=31.37, p<0.0001) and significantly lower at putative centromeres (t=-52.38, p<0.0001) compared to the rest of the genome (Fig 4B). For most regions of the genome, including centromeres and telomeres, d_XY_ was not significantly different among any of the population comparisons (F=0.886, p=0.412). Patterns of sequence divergence varied among population comparisons only in haploblocks corresponding to massive chromosomal inversions (F=152.2, p<0.0001).

In the pairwise comparison of the southernmost populations, the extent of sequence divergence within the haploblocks on chromosomes 18 and 24 (mean d_XY_=0.028) was nearly double the genome-wide average (mean d_XY_=0.015), whereas in the comparisons between northern populations, sequence divergence was lower in the haploblocks on chromosomes 18 and 24 (mean d_XY_=0.009) compared to the rest of the genome (Fig 4C). Furthermore, the inversion haploblock on chromosome 8 showed slightly elevated sequence divergence in the comparison of the southernmost populations (mean d_XY_=0.015) compared to the two other population comparisons (mean d_XY_=0.013), but overall did not show elevated divergence relative to the genome-wide average. The F_ST_ haploblock on chromosome 11 that appeared between NY and NS, showed elevated sequence divergence between those populations (mean d_XY_=0.017) compared to the other population comparisons (mean d_XY_=0.014, Fig 4C).

### Contrasting patterns of divergence and differentiation in inversions and centromeres

Correlations between differentiation (F_ST_) and divergence (d_XY_) revealed distinct patterns across the genome depending on the structural features and levels of gene flow between populations. In genomic windows associated with large chromosomal inversions, high F_ST_ and d_XY_ were observed, particularly between GA and NY where gene flow is extensive (Fig 4D, gold points). These regions with high differentiation and divergence correspond to inversions on chromosomes 18 and 24 for the southernmost population. Elevated differentiation and divergence were also apparent for the inversion on chromosome 11 between NY and NS. Thus, chromosomal inversions, which suppress recombination in a heterozygous state, maintain high levels of both differentiation and divergence between populations under conditions of high gene flow. Conversely, between NS and QU, where gene flow is limited, chromosomal inversions did not show elevated differentiation or sequence divergence. In addition, regions of the genome without inversions exhibited lower correlations between differentiation and divergence, indicating less pronounced divergent selection in those areas in all population comparisons. Notably, in putative centromeres, which are characterized by consistently low recombination, the pattern diverged. Here, high F_ST_ was accompanied by low d_XY_ (Fig 4D, blue points), suggesting these regions are not associated with divergence with gene flow. Instead, in centromeres, where recombination is consistently low, stronger effects of linked selection reduce within-population diversity and drive differentiation between populations, without increasing sequence divergence, indicating that centromeres are unlikely to play a role in preserving locally adapted differences in the face of gene flow.

### Diversity estimates reflect divergent selection in inversions but not centromeres

Patterns of genome-wide π resembled patterns of d_XY_, with similar dramatic oscillations within chromosomes, and dips and peaks corresponding to putative centromeres and telomeres, respectively (Fig S5). Mean genome-wide π varied slightly among populations (F=421.8, p<0.0001), with higher levels in southern populations (GA mean π=0.15, NY mean π=0.16) compared to populations in the north (NS mean π=0.14, QU mean π=0.13). In centromeric regions, π was consistently lower in all populations (Fig 4E). Within chromosomal inversions, π was higher in NY compared to GA, but the same in all other regions of the genome. The inversion on chromosome 11 showed decreased π in the northernmost population, whereas the inversions on chromosomes 8 and 18 showed decreased π in the southernmost population, and for the inversion on chromosome 24, π was lower in NS (Fig 4F). Furthermore, levels of π within populations were positively correlated with levels of d_XY_ and negatively correlated with levels of F_ST_ between populations except at inversions, where differentiation and divergence far exceeded expectations from their levels of diversity (Fig S6). Tajima’s D estimates further supported these patterns. Strongly negative values of Tajima’s D, indicating an excess of rare polymorphisms and possible signals of population expansion or positive selection, were observed in southern populations (GA = −1.2, NY = −1.1). At centromeres, Tajima’s D was lowest in all populations, with northern populations showing negative values despite overall positive Tajima’s D across the genome (NS = 0.2, QU = 0.6), which suggests balancing selection or population contraction (Fig S7).

### Recombination landscapes correlate with diversity, differentiation, and various genome features

We analyzed the relationship between recombination rates and genomic features (Fig S8), focusing on the GA recombination map because the reference genome was assembled using an individual from this population. Our analysis examined genetic diversity within GA and sequence divergence between GA and NY. We observed a positive correlation between recombination rates and nucleotide diversity (τ=0.25, p<0.0001) as well as sequence divergence (τ=0.24, p<0.0001), indicating that regions with higher recombination harbor more genetic variation and greater divergence between these populations. In contrast, recombination rates were negatively correlated with genetic differentiation (τ=-0.10, p<0.0001), suggesting that low-recombination regions are more differentiated. Furthermore, we observed several significant correlations between recombination rates and various genomic features. Recombination rates showed a weak positive correlation with GC content (τ=0.04, p<0.0001) and a similarly weak negative correlation with exon density (τ=-0.05, p<0.0001), indicating that regions with higher recombination tend to have slightly more GC-rich sequences but slightly fewer exons. Additionally, there was a notable positive correlation between recombination rates and tandem repeat content (τ=0.14, p<0.0001), suggesting that regions with more tandem repeats experience higher recombination rates.

## Discussion

Theory predicts that higher levels of gene flow will result in more clustered genetic architectures, with the spatial arrangement of loci underlying adaptive divergence occurring in closer proximity in the genome (16, 63, 64). By examining 168 whole genomes from four populations that span a gradient of gene flow and differentiation, we provide insights into how the interaction between gene flow and selection can drastically shape genome evolution within a species. Our findings indicate that the landscape of adaptive divergence is correlated with patterns of gene flow, with clustering of differentiated regions in the genome intensifying with increasing gene flow between populations, and provide an elegant example supporting theoretical expectations about divergence with gene flow.

The Atlantic silverside is known for its prominent pattern of local adaptation across the latitudinal gradient of its range, which is partitioned into three regional subdivisions with varying levels of gene flow (reviewed in 56, 58). Initial genomic work in this species revealed a dramatic clustering pattern for differentiated loci in the genome in highly connected populations, suggesting that these populations may represent an extreme case of heterogeneity in levels of differentiation across the genome (59). With access to more accurate information about the local genomic landscape, we find that the pattern of genome-wide differentiation between the two southernmost populations is even more striking than it initially appeared. With very high levels of gene flow, genomic differentiation between populations is exclusively constrained to regions of low recombination, resulting in peaks and blocks of differentiation that protrude from an otherwise homogeneous genomic background. This suggests that low-recombination regions are not just favored, but may be essential for divergent selection to persist in the face of high gene flow.

While wide blocks of elevated differentiation coincide with chromosomal inversions that only experience suppressed recombination in a heterozygous state, narrow peaks overlap putative centromeres, where recombination is consistently suppressed. While the differences between centromeres and inversions as recombination modifiers is well known, studies comparing their relative roles in shaping patterns of genomic differentiation are limited. Our results provide an important contribution for our understanding of genome evolution by distinguishing patterns of differentiation and divergence between inversions and centromeres, which despite their fundamental differences, are rarely distinguished in such a way. We discuss the patterns and the likely contributing evolutionary processes first for inversions, then for centromeres.

### Inversion haploblocks show evidence of adaptive divergence with gene flow

Analyzing genome-wide patterns in light of a high-quality reference genome, we confirmed that the massive haploblocks, initially evidenced in transcriptome-level data (59, 65), coincide with segregating chromosomal inversions. As hypothesized based on their level and extent of differentiation, these inversion haploblocks also show elevated sequence divergence, supporting their role in facilitating adaptation with gene flow. The suppression of recombination between alternate arrangements of these inversions explains how isolated blocks of high differentiation are maintained in otherwise largely undifferentiated genomes in the face of gene flow among silverside populations, especially south of NY.

Evolutionary models predict that when an inversion occurs, it leads to a marked reduction in diversity within the two arrangements, resembling a selective sweep or bottleneck, especially if the inversion polymorphism is balanced in the population (66). Over time, this suppression of recombination allows the two arrangements to build up sequence divergence and reintroduce variability through gene flow and new mutations, particularly at increasing distances from the inversion breakpoints (66, 67). Inversions that have persisted for a longer period are expected to show high d_XY_ and a significant number of fixed differences (high F_ST_) between the haplotypes, along with reduced π, and low Tajima’s D values, reflecting the accumulation of genetic differences over time. In contrast, more recent inversions typically display lower d_XY_ and F_ST_, with slight reductions in π and near-neutral Tajima’s D values, indicating that there has been less time for divergence.

Patterns of diversity and differentiation across inversion haploblocks suggest different evolutionary histories. Haploblocks 18 and 24 share characteristics of high sequence divergence, tens of thousands of fixed differences, and low π and Tajima’s D, suggesting they are relatively old. In contrast, haploblock 8, the largest, appears more recent, with fewer fixed differences and slight reductions in π. Haploblock 11, with a dramatic reduction in π at one location, QU, shows recent differentiation, with Tajima’s D suggesting a selective sweep or a recent inversion. Without information that explicitly links adaptive phenotypic variation to patterns of genetic variation in this species, it is not yet possible to discern the relative roles of these multiple inversions in adaptation. The evolutionary history of the Atlantic silverside, shaped by Pleistocene glacial cycles, reveals two waves of post-glaciation colonization from the south, around 16,000 and 8,000 years ago, forming the northern NS and QU populations. The inversions likely evolved south of the glacial front, conferring an adaptive advantage in colder climates and enabling the fish to track receding cold waters as they expanded northward. Future research could further clarify the role of inversions in facilitating adaptation.

### Centromere peaks do not show evidence of adaptive divergence with gene flow

In addition to the large haploblocks discussed above, we identified mountain-like peaks in differentiation (F_ST_) in the south, one for each chromosome, clearly associated with dips in π and d_XY_, and harbored within putative centromeres. In these areas, π and d_XY_ are depressed in all populations, but the corresponding F_ST_ peaks are evident only in the south, where genome-wide differentiation between populations is low. High F_ST_ but low d_XY_ at centromeres relative to the genomic background suggests that in these consistently low-recombination regions, stronger effects of linked selection reduce within-population diversity and drive differentiation between populations, with low sequence divergence indicating an unlikely role for centromeres in preserving locally adapted differences despite gene flow. These patterns offer a clear example of how reduced diversity resulting from areas of low recombination such as centromeres can be mistakenly associated with adaptive divergence in the face of gene flow, especially when a high-quality reference genome and recombination maps are not available. Although the estimates of the recombination landscape and putative centromeric positions are based on reduced-representation sequence data (61), and do not provide the same resolution as the whole genome data, analyzing these data together revealed important insights into the genomic features underlying patterns of differentiation across the genome.

Genomic islands of differentiation, often underpinned by regions of low recombination, have been described in many taxa (12, 68–71), but most studies do not differentiate between conditional low-recombination regions, like inversions, and consistent ones, such as centromeres. The centromeric patterns we observed here are more dramatic and consistent across the genome than what has been shown in other species, particularly given our focus on intraspecific populations. While the patterns may be due to extremely high levels of gene flow and/or strong selection acting on these populations, they could also be due to highly polygenic trait architectures. Determining whether these centromeric regions harbor loci contributing to adaptation is an important but challenging next step, and may be better characterized in the future with more long-read sequencing and fine-scale trait mapping. Additionally, centromere drive, a form of meiotic drive that occurs during female meiosis, may be a possible explanation for the consistent peaks in F_ST_ observed on every chromosome. According to the centromere drive hypothesis, a centromere can be retained in a female gamete (i.e., in the oocyte rather than the polar body) more often during meiosis, and can therefore act like a selfish genetic element driving non-Mendelian segregation (reviewed in 72, 73). This usually results in fitness costs and genetic conflict in the genome that imposes strong selective pressures on centromeric DNA. In populations that become isolated, the competition between centromere sequences can quickly drive differentiation at these regions. For instance, in medaka (74) and pink salmon (75), centromeric differences are thought to play a role in speciation. Further studies examining segregation distortion in crosses are needed to test for the potential role of centromere drive in shaping genome evolution in Atlantic silversides.

## Conclusion

Our study provides a comprehensive analysis of how gene flow, selection, and recombination interact to shape patterns of genomic differentiation within the Atlantic silverside. By distinguishing between the contributions of inversions and centromeres, we have uncovered the complex genomic landscape that underlies adaptive divergence in this species. The clustering of differentiated regions in response to high gene flow underscores the critical role of genomic architecture in facilitating adaptation. Our findings not only confirm theoretical predictions about divergence with gene flow but also offer new insights into the relative influence of inversions and centromeres in maintaining genetic differentiation. As we continue to unravel the genetic basis of adaptation, further investigation into the functional roles of these genomic regions will be crucial for understanding the mechanisms driving speciation and adaptation in high gene flow environments.

## Materials and Methods

### Whole genome resequencing and variant calling

To optimally explore the role of genome structure in the distribution and levels of diversity and differentiation, we used the Atlantic silverside reference genome v2 (76), which was improved by anchoring the first version of the reference genome (60) to a species-specific linkage map (61). We then re-examined low-coverage whole genome resequencing data (59) for 42-50 wild-caught Atlantic silverside individuals from four locations: Jekyll Island, Georgia (GA), Patchogue, New York (NY), Minas Basin, Nova Scotia (NS), and Magdalen Island, Quebec (QU) (Fig 1). To avoid potential bias due to variation in sample sizes, we only included 42 individuals from each population, removing individuals with the least amount of data.

Adapters were trimmed from sequence reads using Trimmomatic v.0.36 with seed matches=2, palindrome clip threshold=30, simple clip threshold=10, and minAdapterLength=4 (77). Paired, adapter-clipped reads were then mapped to the reference genome using Bowtie2 v.2.2.9 (78) with the --very-sensitive preset option. Reads with mapping qualities below 20 were filtered out and the remaining reads sorted using Samtools v.1.9 (79). Alignment bam files from each lane were then merged for each individual. We then removed duplicated reads using MarkDuplicates v.2.9 from Picard tools (broadinstitute.github.io/picard) and realigned reads around indels using IndelRealigner from GATK (80).

To account for the uncertainty about individual genotypes associated with low-coverage data, we used ANGSD v.0931 (81) and conducted our population genomics analyses within a probabilistic framework based on genotype likelihoods. We first examined the sequencing depth distribution across all individuals with the commands -doCounts and -doDepth in ANGSD, using the mode ± 2 standard deviations to establish minimum and maximum depth filters for calling SNPs. We used all individuals to call SNPs globally (p-value=10^−5^), considering only sites with a minimum combined sequencing depth of 120, maximum combined sequencing depth of 428, mapped reads from at least half of the individuals (n=84), and removed sites with a global minor allele frequency below 0.01. Then, we supplied a list of global SNPs using the -sites option to estimate allele frequencies in each of the four populations separately, excluding sites in the tail ends of the depth distributions for each population. Sites with read depth less than 20 were excluded from each population, sites with read depth more than 150 were excluded for NY, and sites with read depth more than 120 were excluded for all other populations.

### Estimating differentiation, diversity and linkage disequilibrium

To investigate population structure, we first conducted a principal components analysis (PCA) by computing eigenvectors in R from the covariance matrix between individuals estimated in PCAngsd (82). We estimated population genetic parameters in non-overlapping 50 kb windows. To ensure we compared the same non-overlapping 50 kb windows across analyses, we calculated the window intervals with the command *makewindows* from BEDTools v.2.29.2 (83) based on the lengths of chromosomes of the reference genome. To calculate pairwise genetic differentiation (F_ST_) between populations, we generated the joint SFS (2dSFS) for each pair of neighboring populations from their respective site frequency spectra using realSFS *fst stats*. We computed weighted Weir and Cockerham F_ST_ averages as the ratio of the sum of alpha (between-population variance) (84) to the sum of alpha plus beta (within-population variance) across all sites in 50 kb windows. We also calculated the pairwise absolute allele frequency differences (AFD) between neighboring populations using the minor allele frequency estimates for each population as an alternative to F_ST_ that is more sensitive to weak population differentiation (62).

To calculate nucleotide diversity (π) and Tajima’s D within populations and pairwise sequence divergence (d_XY_) between populations, we used the SFS based on all sites (i.e., variant and invariant sites, so no SNP calling, only the depth filter) as a prior. We estimated per-site thetas (population scaled mutation rate) for each population with -doThetas then used thetaStat to calculate π and Tajima’s D averages per 50 kb window, filtering out windows with fewer than 100 sites for estimates of π and Tajima’s D. To calculate d_XY_, we used a custom python script *dxy_wsfs.py* (85) after estimating the 2dSFS in ANGSD using all sites for each window for each pair of neighboring populations.

### Characterizing genome features

We obtained pedigree-based recombination rates in cM/Mb from Akopyan et al. (61), including three recombination maps for NY, GA, and an interpopulation cross. Due to drastic differences in recombination between sexes (i.e. heterochiasmy) observed in this species, with male recombination restricted to the terminal ends of chromosomes (61), we focused our analysis on female recombination rates, which we averaged into the same 50 kb windows described above. Because the pedigree-based recombination information was based on reduced representation sequencing data, the amount of data missing in windows was relatively high, with 46%, 52%, and 35% of 50 kb windows missing data on recombination rates for GA, NY, and hybrid maps, respectively.

To evaluate genome-wide associations between recombination rates and genomic features that are known to correlate with recombination landscapes in other species, we characterized gene density and GC content, as recombination events tend to localize in GC-rich and/or gene-dense regions in other vertebrate species (86–88). To identify coding regions in the silverside genome, we obtained annotation coordinates (76) and calculated the total number and average proportion of exons and coding sequences (including exons and the 5’ and 3’ UTRs) for each 50 kb window with BEDTools *intersect*. We calculated GC content by obtaining the base composition of each window of the reference genome using BEDTools *nuc.* Data summaries were performed using the tidyverse package in R v. 4.0.0 (89).

We also identified the location of known segregating inversions and putative centromeres and telomeres, which are typically associated with recombination cold- and hot-spots (90, 91). We obtained and lifted over inversion positions from Akopyan et al. (61), then estimated the putative locations of centromeres and telomeres using a combination of approaches. First, we used the three recombination maps to identify heterochromatin boundaries typical of centromeres and telomeres based on patterns of marker density and distribution, and local changes in recombination rates with BREC (92). Second, we used Tandem Repeats Finder v.4.09 (93) to identify repeats in the reference genome with pattern size less than 500 bp that are generally associated with telomeres in eukaryotes. We used trfparser v.1 (trfparser.sourceforge.io/) to parse the output, then filtered for repeats based on the telomere repeat motif (TTAGGG) that is conserved among vertebrates (94) to refine our estimates of telomere positions. We used BEDTools intersect to categorize each 50kb window as an inversion, a telomere, a centromere, or none of those. Windows falling into more than one category were assigned to the category with more support, prioritizing inversions, followed by telomeres, then centromeres.

### Evaluating landscapes of diversity, differentiation, and recombination across genomic features

We plotted all estimates of population statistics and genome parameters in 50 kb windows to visualize genome-wide patterns using Manhattan plots. We analyzed the genomic distribution of F_ST_ outlier loci across chromosomes to assess clustering patterns. For each of the three pairwise comparisons of neighboring populations, we identified outliers as loci with F_ST_ values exceeding three standard deviations above the mean. Using a permutation test, we randomly redistributed the outliers across chromosomes, weighted by chromosome size, to generate a null distribution of expected variance in outlier counts. This process was repeated 10,000 times to establish a baseline for random clustering. We compared the observed variance in outlier distribution against this null distribution to determine the non-randomness of the clustering in each comparison.

To compare how population genetic estimates varied among chromosomal regions and among pairwise comparisons of populations, we used linear mixed models with population comparison and chromosomal region as random effects, then used least-square means for post hoc pairwise comparisons. We also evaluated consistency between relative and absolute measures of differentiation by correlating windowed estimates of F_ST_ and d_XY_ across pairs using a Kendall’s rank correlation test. The statistical analyses were conducted in R v. 4.0.0 (89).

## Acknowledgements

We would like to thank David Conover for providing access to the silverside samples analyzed in this study, and Harmony Borchardt-Wier and Cornell’s Biotechnology Resource Center for help with library preparation for low-coverage whole genome sequencing. This study was funded through a National Science Foundation grant to NOT (OCE-1756316).

## Author Contributions

M.A., A.T., A.J., A.P.W. and N.O.T. designed the study. A.J. mapped the population data to the reference genome, and M.A. conducted the data analysis. M.A. drafted the manuscript with critical input from all authors.

## Supplementary Figures

**Figure S1.**
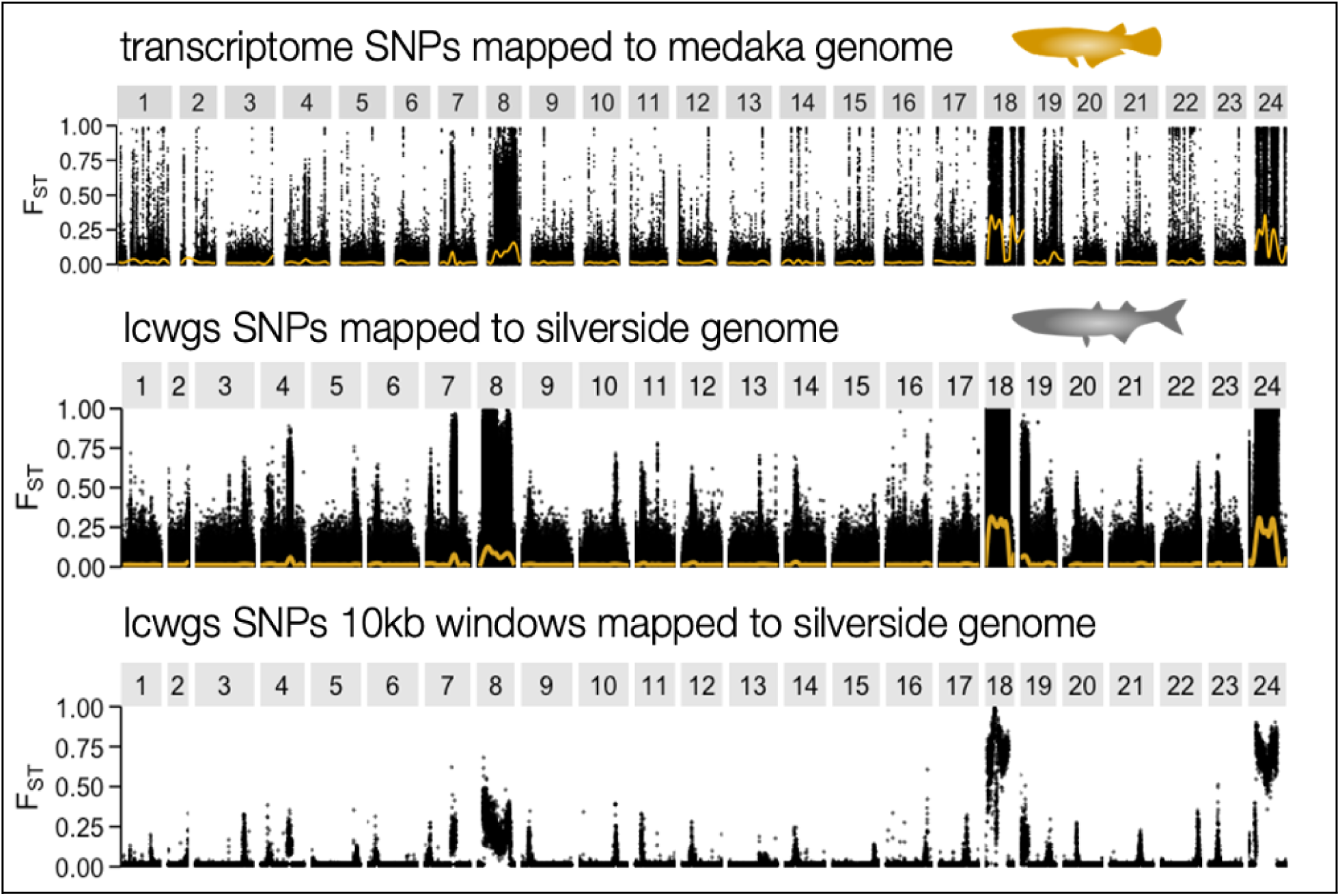
Manhattan plots of F_ST_ between GA and NY for (top) individual SNPs ordered along Atlantic silverside transcriptome contigs anchored to the reference genome for the medaka (from Wilder et al. 2020), (center) individual SNPs ordered along chromosomes in the Atlantic silverside genome (this study), and (bottom) averaged within 50 kb windows along the Atlantic silverside genome (this study).

**Figure S2.**
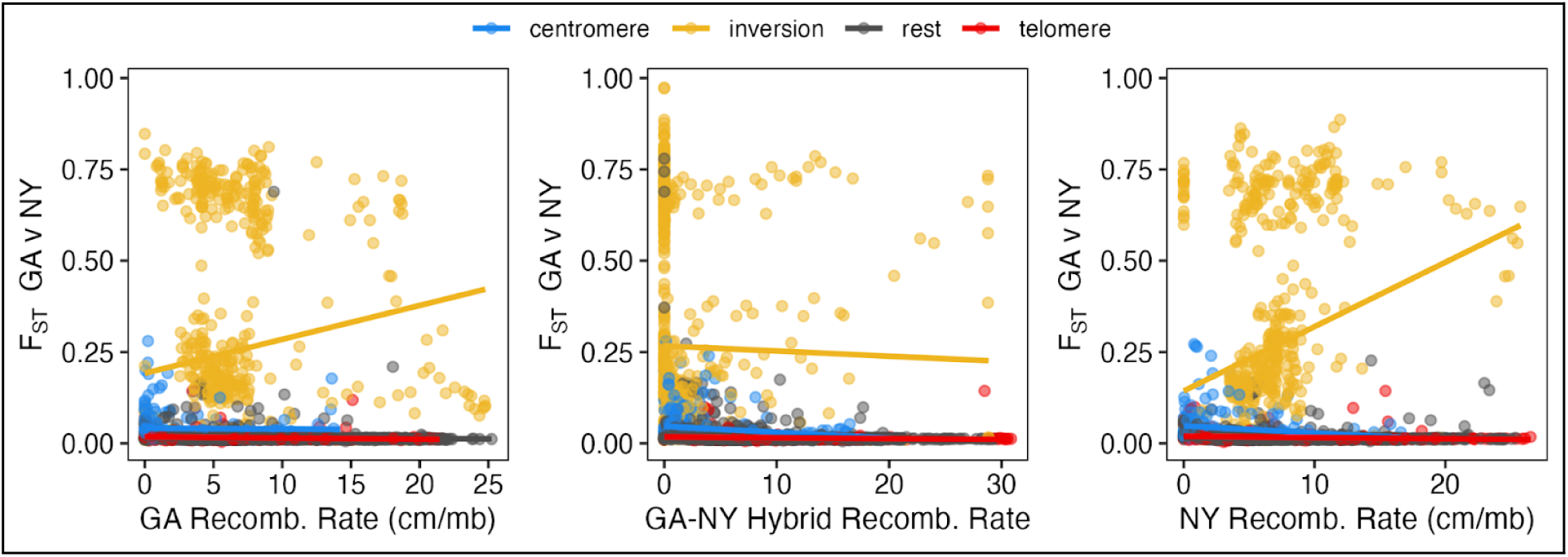
Differentiation (F_ST_) between GA and NY populations correlates with recombination rates across genomic regions. Scatter plots show the relationship between F_ST_ (GA vs NY) and recombination rates using the GA recombination map (left), GA-NY hybrid recombination map (center), and NY recombination map (right). Solid lines represent linear regression fits. Inversions exhibit a positive correlation between recombination rates and F_ST_ in both the GA map (τ = 0.18, p < 0.001) and the NY map (τ = 0.17, p < 0.001), but a negative correlation (τ = −0.08, p = 0.004) in the GA-NY hybrid recombination map. All other genomic regions show a negative correlation across all maps.

**Figure S3.**
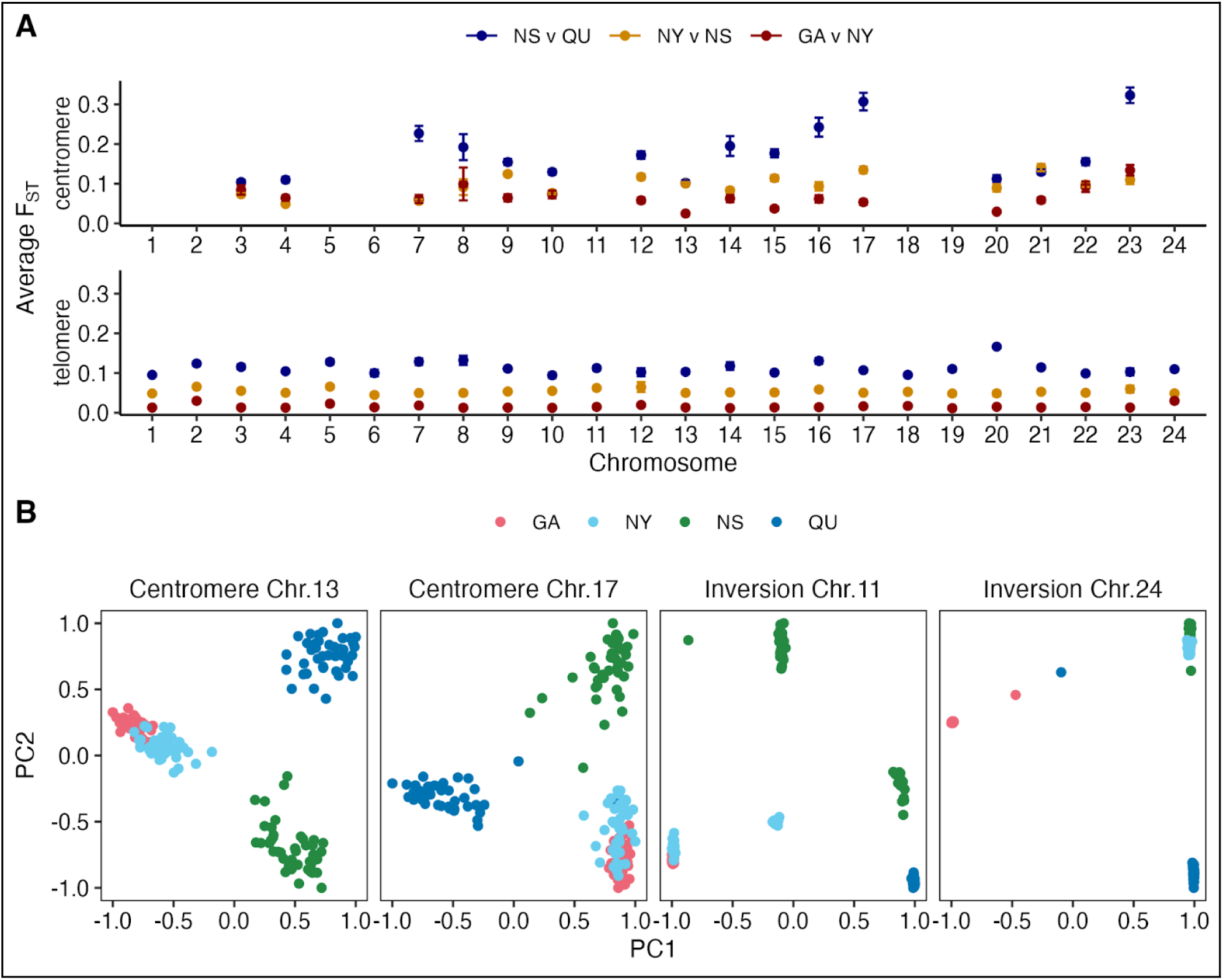
A) Average F_ST_ per chromosome between neighboring populations for SNPs located within putative centromere regions (top panel) and SNPs located within putative telomeres (bottom panel). Data are missing for chromosomes where putative centromeres could not be localized. Note the much greater consistency of patterns across chromosomes for the telomeres compared to the centromeres. B) PCA based on SNPs in centromeric regions of chromosomes 13 and 17 and based on inverted regions of chromosomes 11 and 24. Inversions show a typical pattern of three main clusters that represent the two inversion homozygotes and the heterozygotes, whereas centromeric loci show a similar pattern but with more variation as evidenced by the greater spread of individuals.

**Figure S4.**
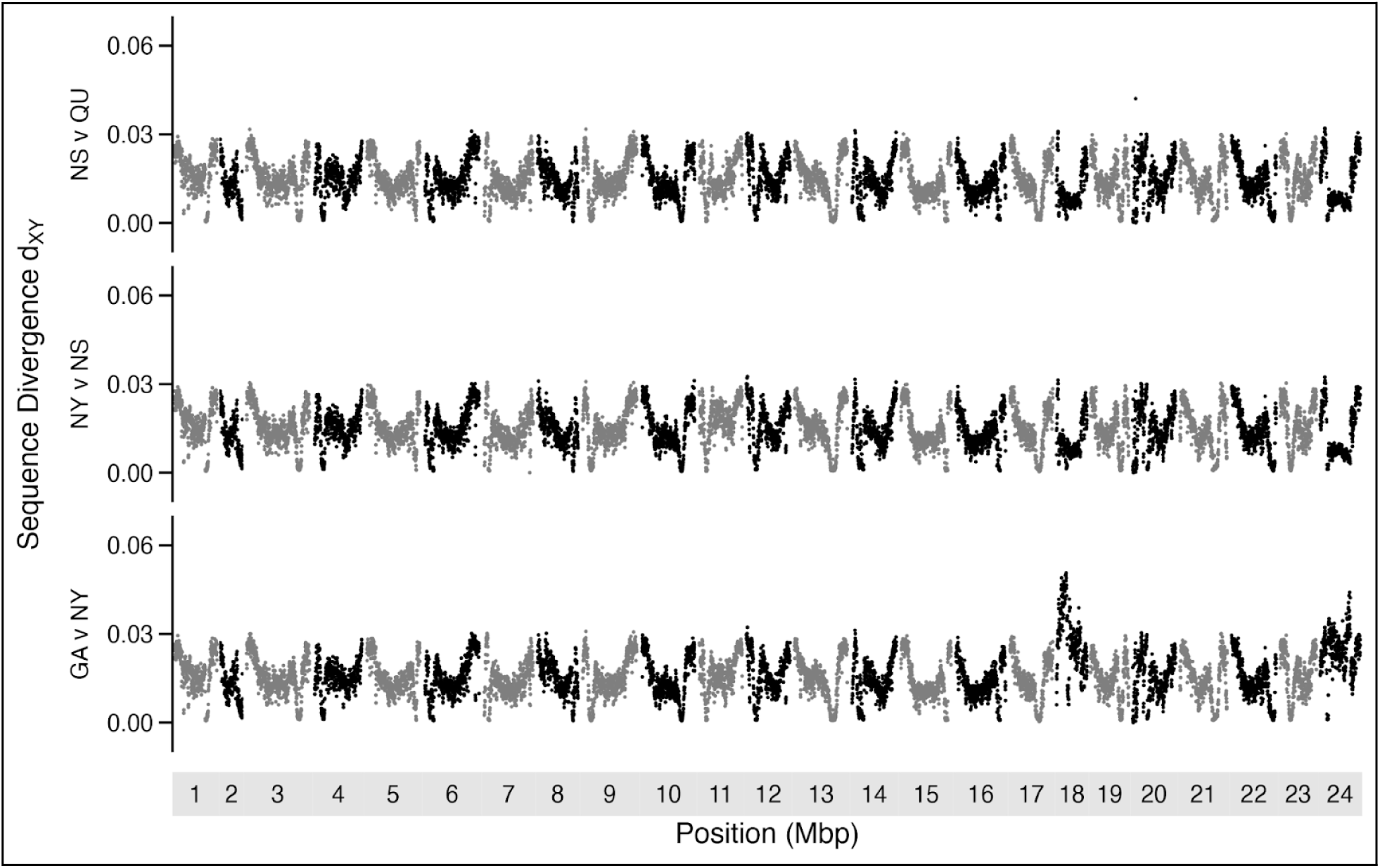
Manhattan plots showing absolute divergence (d_XY_) between neighboring populations (comparisons arranged from north to south) averaged in 50 kb windows ordered along chromosomes in the Atlantic silverside genome.

**Figure S5.**
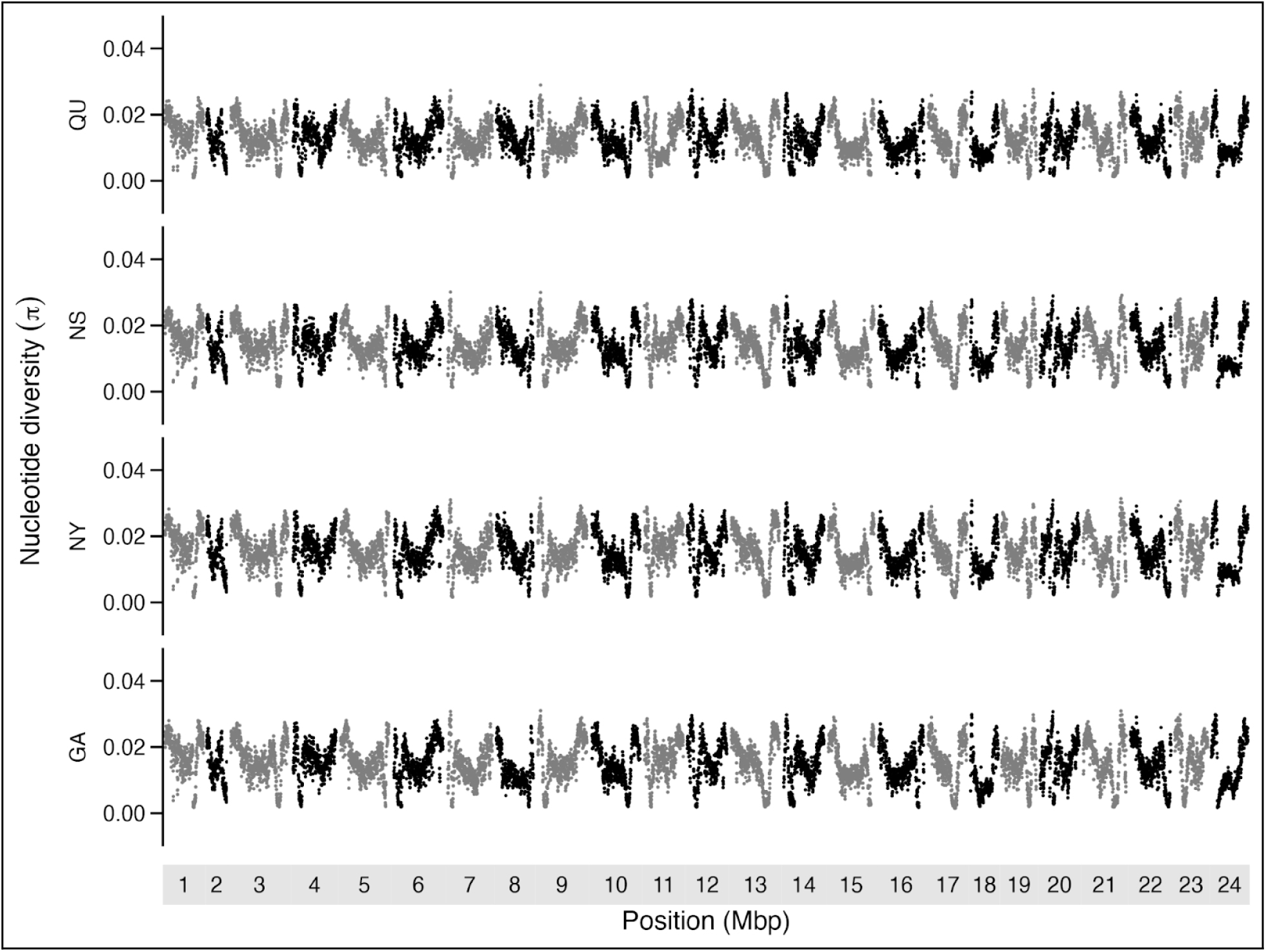
Manhattan plots of nucleotide diversity (π) within each of the four populations (arranged from north to south from top to bottom) averaged within 50 kb windows ordered along chromosomes in the Atlantic silverside genome.

**Figure S6.**
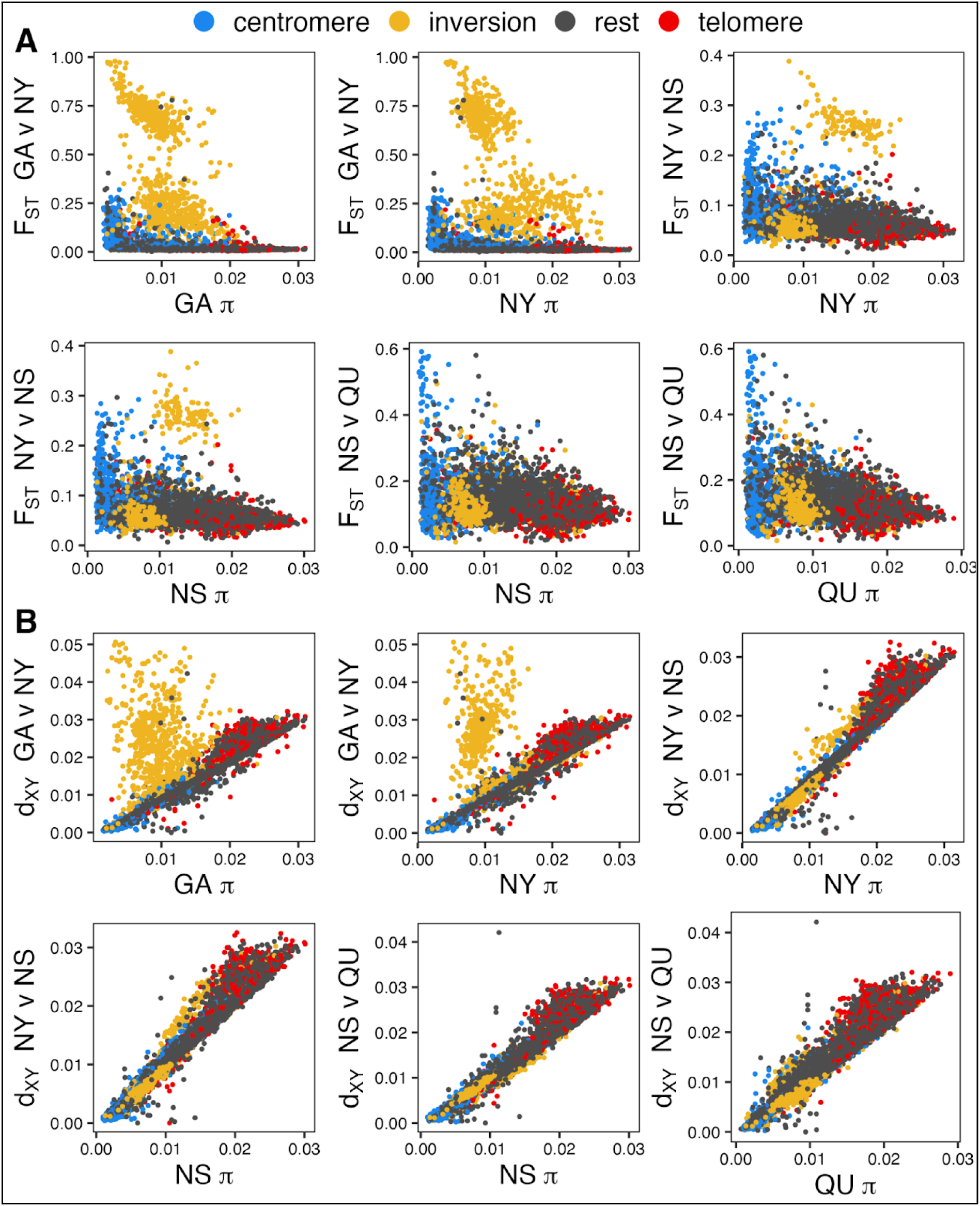
Correlation between nucleotide diversity (π) and genetic differentiation (F_ST_ and d_XY_) across genomic regions. Panel A shows the relationship between diversity (π) on the x-axis and differentiation (F_ST_) on the y-axis for neighboring population comparisons. Panel B displays the relationship between diversity (π) and sequence divergence (d_XY_) on the y-axis. Each point represents the average value within 50 kb windows, colored by genomic region.

**Figure S7.**
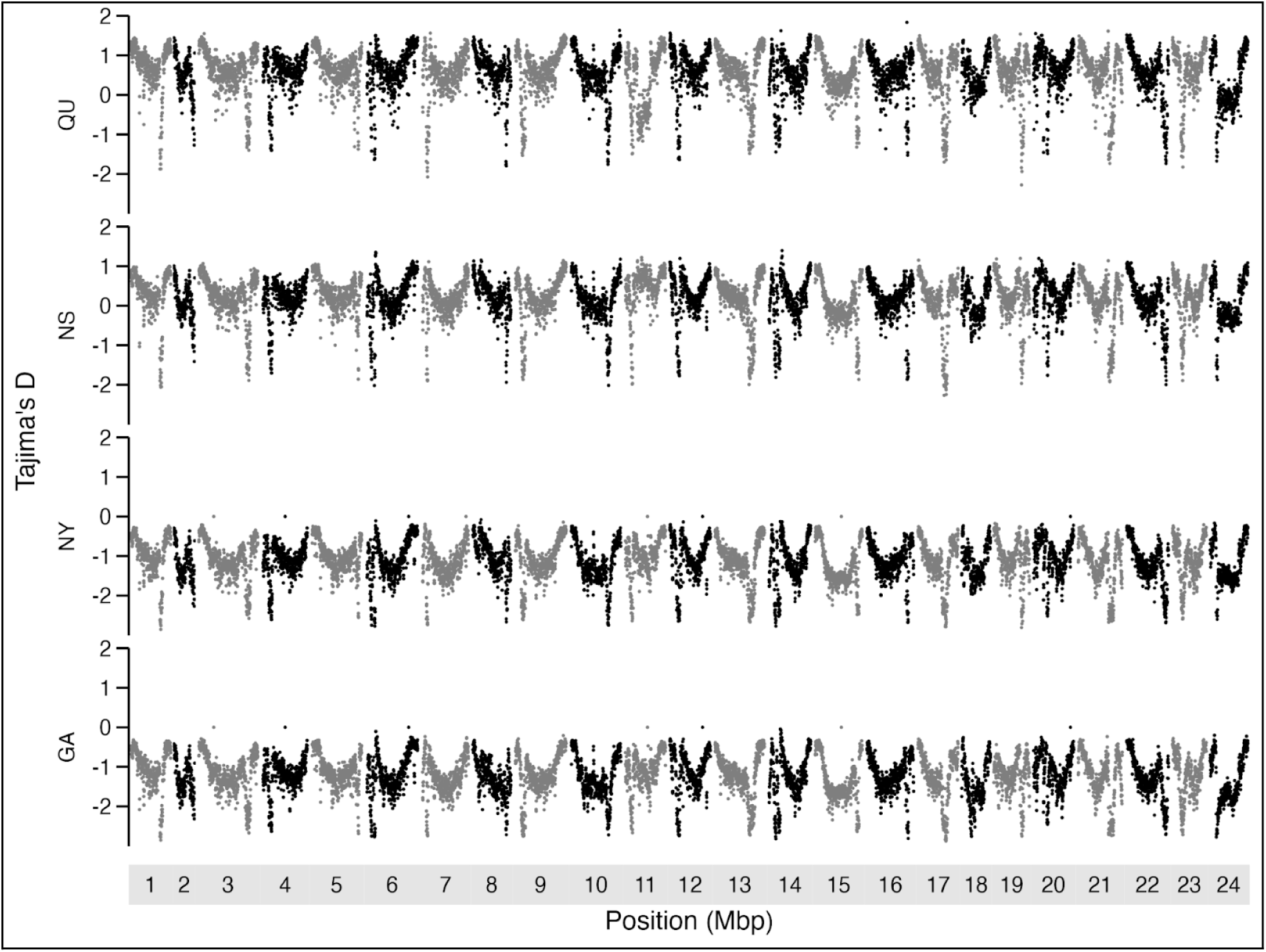
Manhattan plots of Tajima’s D within each of the four populations (arranged from north to south) averaged within 50 kb windows ordered along chromosomes in the Atlantic silverside genome.

**Figure S8.**
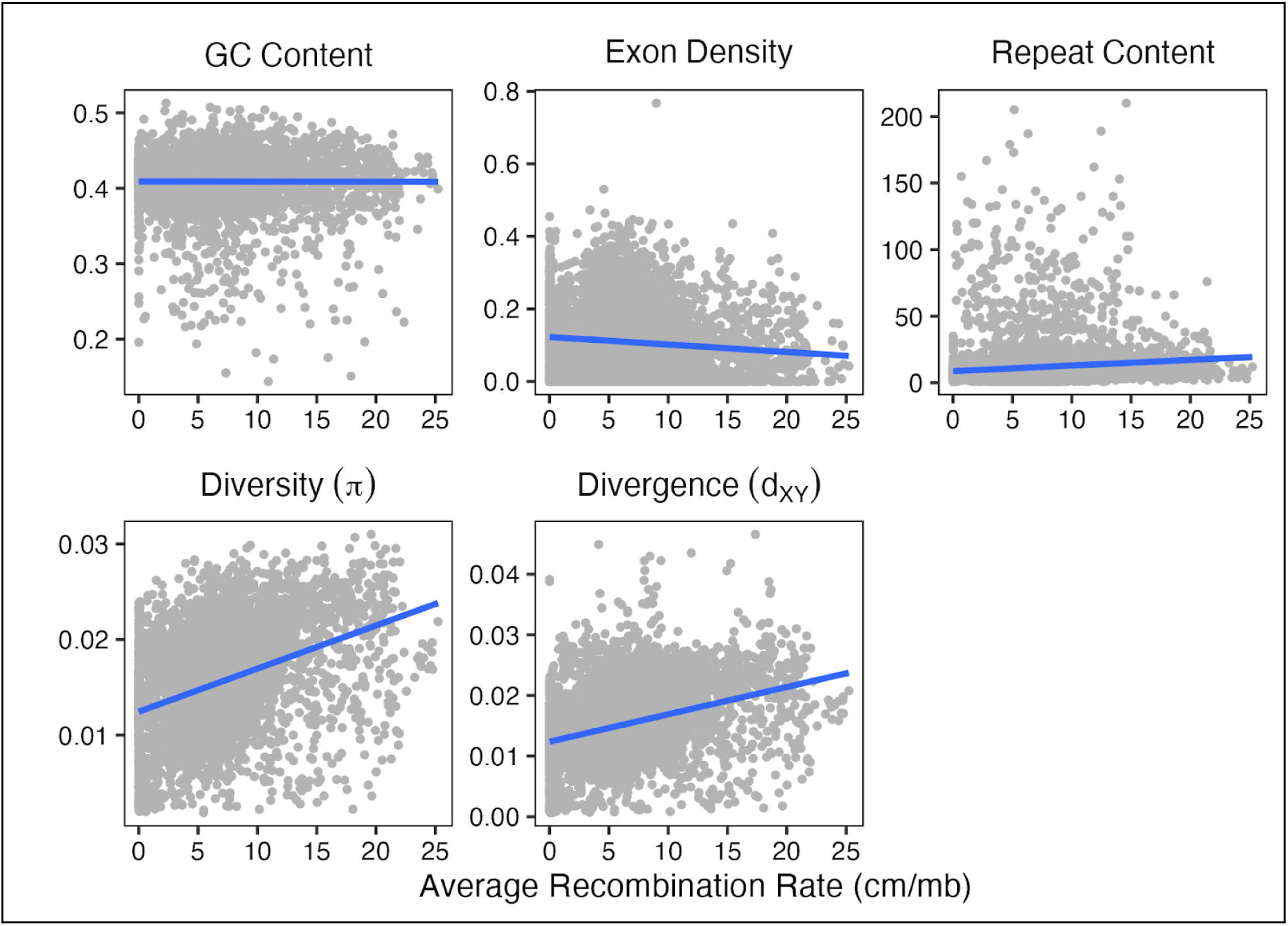
Scatter plots illustrate the relationships between average recombination rates (cM/Mb) in the GA genome and various genomic features: GC content, exon density, repeat content (top) and nucleotide diversity within GA populations and sequence divergence between GA and NY populations (bottom). Each point represents a genomic region, and solid lines indicate linear regression fits. Significant positive correlations are observed between recombination rates and nucleotide diversity (τ = 0.25, p < 0.0001), sequence divergence (τ = 0.24, p < 0.0001), and repeat content (τ = 0.14, p < 0.0001). Recombination rates also show a weak positive correlation with GC content (τ = 0.04, p < 0.0001) and a weak negative correlation with exon density (τ = –0.05, p < 0.0001).

## References

1. M. Slatkin, Gene flow and the geographic structure of natural populations. Science 236, 787–792 (1987).

2. G. García-Ramos, M. Kirkpatrick, Genetic models of adaptation and gene flow in peripheral populations. Evolution 51, 21–28 (1997).

3. J. Felsenstein, The theoretical population genetics of variable selection and migration. Annu. Rev. Genet. 10, 253–280 (1976).

4. T. Lenormand, Gene flow and the limits to natural selection. Trends Ecol. Evol. 17, 183–189 (2002).

5. S. Gavrilets, Fitness Landscapes and the Origin of Species (MPB-41) (Princeton University Press, 2004).

6. A. Tigano, V. L. Friesen, Genomics of local adaptation with gene flow. Mol Ecol 25, 2144–2164 (2016).

7. C. Mérot, R. A. Oomen, A. Tigano, M. Wellenreuther, A roadmap for understanding the evolutionary significance of structural genomic variation. Trends Ecol. Evol. 35, 561–572 (2020).

8. K. Lucek, Z. Gompert, P. Nosil, The role of structural genomic variants in population differentiation and ecotype formation in *Timema cristinae* walking sticks. Mol. Ecol. 28, 1224–1237 (2019).

9. M. H. Weissensteiner, et al., Discovery and population genomics of structural variation in a songbird genus. Nat. Commun. 11, 3403 (2020).

10. T. Hämälä, et al., Genomic structural variants constrain and facilitate adaptation in natural populations of *Theobroma cacao*, the chocolate tree. Proc. Natl. Acad. Sci. 118, e2102914118 (2021).

11. D. Ortiz-Barrientos, M. E. James, *Evolution* of recombination rates and the genomic landscape of speciation. J. Evol. Biol. 30, 1519–1521 (2017).

12. K. Samuk, et al., Gene flow and selection interact to promote adaptive divergence in regions of low recombination. Mol. Ecol. 26, 4378–4390 (2017).

13. S. H. Martin, J. W. Davey, C. Salazar, C. D. Jiggins, Recombination rate variation shapes barriers to introgression across butterfly genomes. PLOS Biol. 17, e2006288 (2019).

14. M. Kirkpatrick, N. Barton, Chromosome inversions, local adaptation and speciation. Genetics 173, 419–434 (2006).

15. A. A. Hoffmann, L. H. Rieseberg, Revisiting the impact of inversions in evolution: From population genetic markers to drivers of adaptive shifts and speciation? Annu. Rev. Ecol. Evol. Syst. 39, 21–42 (2008).

16. S. Yeaman, Genomic rearrangements and the evolution of clusters of locally adaptive loci. Proc. Natl. Acad. Sci. 110, E1743–E1751 (2013).

17. M. Wellenreuther, L. Bernatchez, Eco-evolutionary genomics of chromosomal inversions. Trends Ecol Evol (2018).

18. A. H. Sturtevant, G. W. Beadle, The relations of inversions in the X chromosome of Drosophila melanogaster to crossing over and disjunction. Genetics 21, 554–604 (1936).

19. A. Navarro, E. Betrán, A. Barbadilla, A. Ruiz, Recombination and gene flux caused by gene conversion and crossing over in inversion heterokaryotypes. Genetics 146, 695–709 (1997).

20. L. H. Rieseberg, Chromosomal rearrangements and speciation. Trends Ecol Evol 16, 351–358 (2001).

21. M. J. Christmas, et al., Genetic barriers to historical gene flow between cryptic species of alpine bumblebees revealed by comparative population genomics. Mol. Biol. Evol. 38, 3126–3143 (2021).

22. Q. Haenel, T. G. Laurentino, M. Roesti, D. Berner, Meta-analysis of chromosome-scale crossover rate variation in eukaryotes and its significance to evolutionary genomics. Mol Ecol 27, 2477–2497 (2018).

23. G. R. Smith, M. Nambiar, New solutions to old problems: Molecular mechanisms of meiotic crossover control. Trends Genet. 36, 337–346 (2020).

24. Y. Xu, J. Du, Young but not relatively old retrotransposons are preferentially located in gene-rich euchromatic regions in tomato (*Solanum lycopersicum)* plants. Plant J. 80, 582–591 (2014).

25. A. L. Dapper, B. A. Payseur, Connecting theory and data to understand recombination rate evolution. Philos. Trans. R. Soc. B 372, 20160469 (2017).

26. J. Stapley, P. G. D. Feulner, S. E. Johnston, A. W. Santure, C. M. Smadja, Variation in recombination frequency and distribution across eukaryotes: patterns and processes. Philos. Trans. R. Soc. B Biol. Sci. 372, 20160455 (2017).

27. M. Roesti, D. Moser, D. Berner, Recombination in the threespine stickleback genome—patterns and consequences. Mol. Ecol. 22, 3014–3027 (2013).

28. L. Smeds, C. F. Mugal, A. Qvarnström, H. Ellegren, High-resolution mapping of crossover and non-crossover recombination events by whole-genome re-sequencing of an avian pedigree. PLoS Genet. 12, e1006044 (2016).

29. N. Galtier, G. Piganeau, D. Mouchiroud, L. Duret, GC-content evolution in mammalian genomes: The biased gene conversion hypothesis. Genetics 159, 907–911 (2001).

30. M. Rousselle, A. Laverré, E. Figuet, B. Nabholz, N. Galtier, Influence of recombination and GC-biased gene conversion on the adaptive and non-adaptive substitution rate in mammals vs. birds. Mol. Biol. Evol. 36, msy243 (2018).

31. C. F. Mugal, C. C. Weber, H. Ellegren, GC-biased gene conversion links the recombination landscape and demography to genomic base composition. BioEssays 37, 1317–1326 (2015).

32. K. Paigen, et al., The recombinational anatomy of a mouse chromosome. PLoS Genet. 4, e1000119 (2008).

33. B. S. Gaut, S. I. Wright, C. Rizzon, J. Dvorak, L. K. Anderson, Recombination: an underappreciated factor in the evolution of plant genomes. Nat. Rev. Genet. 8, 77–84 (2007).

34. G. P. Tiley, J. G. Burleigh, G. Burleigh, The relationship of recombination rate, genome structure, and patterns of molecular evolution across angiosperms. BMC Evol. Biol. 15, 194 (2015).

35. A. Wallberg, S. Glémin, M. T. Webster, Extreme recombination frequencies shape genome variation and evolution in the honeybee, *Apis mellifera*. PLoS Genet. 11, e1005189 (2015).

36. J. C. Jones, A. Wallberg, M. J. Christmas, K. M. Kapheim, M. T. Webster, Extreme differences in recombination rate between the genomes of a solitary and a social bee. Mol. Biol. Evol. 36, 2277–2291 (2019).

37. A. P. i Torres, et al., The fine-scale recombination rate variation and associations with genomic features in a butterfly. Genome Res. 33, 810–823 (2023).

38. G. A. T. McVean, et al., The fine-scale structure of recombination rate variation in the human genome. Science 304, 581–584 (2004).

39. M. A. F. Noor, S. M. Bennett, Islands of speciation or mirages in the desert? Examining the role of restricted recombination in maintaining species. Heredity 103, 439–444 (2009).

40. T. E. Cruickshank, M. W. Hahn, Reanalysis suggests that genomic islands of speciation are due to reduced diversity, not reduced gene flow. Mol. Ecol. 23, 3133–3157 (2014).

41. K. E. Lotterhos, The effect of neutral recombination variation on genome scans for selection. G3: Genes, Genomes, Genet. 9, 1851–1867 (2019).

42. T. R. Booker, S. Yeaman, M. C. Whitlock, Variation in recombination rate affects detection of outliers in genome scans under neutrality. Mol. Ecol. 29, 4274–4279 (2020).

43. R. Burri, et al., Linked selection and recombination rate variation drive the evolution of the genomic landscape of differentiation across the speciation continuum of *Ficedula* flycatchers. Genome Res. 25, 1656–1665 (2015).

44. K. E. Delmore, et al., Comparative analysis examining patterns of genomic differentiation across multiple episodes of population divergence in birds. Evol. Lett. 2, 76–87 (2018).

45. D. E. Irwin, M. Alcaide, K. E. Delmore, J. H. Irwin, G. L. Owens, Recurrent selection explains parallel evolution of genomic regions of high relative but low absolute differentiation in a ring species. Mol. Ecol. 25, 4488–4507 (2016).

46. H. A. Hejase, et al., Genomic islands of differentiation in a rapid avian radiation have been driven by recent selective sweeps. Proc. Natl. Acad. Sci. 117, 30554–30565 (2020).

47. D. A. DeRaad, J. E. McCormack, N. Chen, A. T. Peterson, R. G. Moyle, Combining species delimitation, species trees, and tests for gene flow clarifies complex speciation in scrub-jays. Syst. Biol. 71, 1453–1470 (2022).

48. S. M. Flaxman, A. C. Wacholder, J. L. Feder, P. Nosil, Theoretical models of the influence of genomic architecture on the dynamics of speciation. Mol. Ecol. 23, 4074–4088 (2014).

49. J. Gutiérrez-Valencia, P. W. Hughes, E. L. Berdan, T. Slotte, The genomic architecture and evolutionary fates of supergenes. Genome Biol. Evol. 13, evab057 (2021).

50. S. M. Schaal, B. C. Haller, K. E. Lotterhos, Inversion invasions: when the genetic basis of local adaptation is concentrated within inversions in the face of gene flow. Philos. Trans. R. Soc. B 377, 20210200 (2022).

51. L. A. Hice, T. A. Duffy, S. B. Munch, D. O. Conover, Spatial scale and divergent patterns of variation in adapted traits in the ocean. Ecol. Lett. 15, 568–575 (2012).

52. D. O. Conover, T. M. C. Present, Countergradient variation in growth rate: compensation for length of the growing season among Atlantic silversides from different latitudes. Oecologia 83, 316–324 (1990).

53. S. A. Arnott, S. Chiba, D. O. Conover, Evolution of intrinsic growth rate: metabolic costs drive trade-offs between growth and swimming performance in *Menidia menidia*. Evolution 60, 1269–1278 (2006).

54. J. M. Billerbeck, T. E. Lankford, D. O. Conover, Evolution of intrinsic growth and energy acquisition rates. I. Trade-offs with swimming performance in Menidia menidia. Evolution 55, 1863–1872 (2001).

55. S. B. Munch, D. O. Conover, Rapid growth results in increased susceptibility to predation in *Menidia menidia*. Evolution 57, 2119–2127 (2003).

56. D. O. Conover, S. A. Arnott, M. R. Walsh, S. B. Munch, Darwinian fishery science: lessons from the Atlantic silverside (*Menidia menidia*). Can. J. Fish Aquat. Sci. 62, 730–737 (2005).

57. L. Clarke, B. Walther, S. Munch, S. Thorrold, D. Conover, Chemical signatures in the otoliths of a coastal marine fish, *Menidia menidia*, from the northeastern United States: spatial and temporal differences. Mar. Ecol. Prog. Ser. 384, 261–271 (2009).

58. R. N. Lou, et al., Full mitochondrial genome sequences reveal new insights about post-glacial expansion and regional phylogeographic structure in the Atlantic silverside (*Menidia menidia*). Mar. Biol. 165, 124 (2018).

59. A. P. Wilder, S. R. Palumbi, D. O. Conover, N. O. Therkildsen, Footprints of local adaptation span hundreds of linked genes in the Atlantic silverside genome. Evol. Lett. 4, 430–443 (2020).

60. A. Tigano, et al., Chromosome-level assembly of the Atlantic silverside genome reveals extreme levels of sequence diversity and structural genetic variation. Genome Biol. Evol. 13, evab098- (2021).

61. M. Akopyan, et al., Comparative linkage mapping uncovers recombination suppression across massive chromosomal inversions associated with local adaptation in Atlantic silversides. Mol. Ecol. (2022).

62. D. Berner, Allele frequency difference AFD–An intuitive alternative to F_ST_ for quantifying genetic population differentiation. Genes 10, 308 (2019).

63. S. Via, Divergence hitchhiking and the spread of genomic isolation during ecological speciation-with-gene-flow. Philos. Trans. R. Soc. B Biol. Sci. 367, 451–460 (2012).

64. S. Yeaman, M. C. Whitlock, The genetic architecture of adaptation under migration-selection balance. Evolution 65, 1897–1911 (2011).

65. N. O. Therkildsen, et al., Contrasting genomic shifts underlie parallel phenotypic evolution in response to fishing. Science 365, 487–490 (2019).

66. A. Navarro, A. Barbadilla, A. Ruiz, Effect of inversion polymorphism on the neutral nucleotide variability of linked chromosomal regions in *Drosophila*. Genetics 155, 685–698 (2000).

67. P. Andolfatto, F. Depaulis, A. Navarro, Inversion polymorphisms and nucleotide variability in *Drosophila*. Genet. Res. 77, 1–8 (2001).

68. Y. Shi, B. Zhou, Y. Liang, B. Wang, Linked selection and recombination rate generate both shared and lineage-specific genomic islands of divergence in two independent *Quercus* species pairs. J. Syst. Evol. 62, 505–519 (2024).

69. R. A. Bay, K. Ruegg, Genomic islands of divergence or opportunities for introgression? Proc. R. Soc. B: Biol. Sci. 284, 20162414 (2017).

70. M. Tine, et al., European sea bass genome and its variation provide insights into adaptation to euryhalinity and speciation. Nat. Commun. 5, 5770 (2014).

71. D. Zhang, et al., Genomic differentiation and patterns of gene flow between two long-tailed tit species (*Aegithalos*). Mol. Ecol. 26, 6654–6665 (2017).

72. S. Henikoff, K. Ahmad, H. S. Malik, The centromere paradox: Stable inheritance with rapidly evolving DNA. Science 293, 1098–1102 (2001).

73. M. A. Lampson, B. E. Black, Cellular and molecular mechanisms of centromere drive. Cold Spring Harb. Symp. Quant. Biol. 82, 249–257 (2017).

74. K. Ichikawa, et al., Centromere evolution and CpG methylation during vertebrate speciation. Nat. Commun. 8, 1833 (2017).

75. K. A. Christensen, et al., The pink salmon genome: Uncovering the genomic consequences of a two-year life cycle. PLoS ONE 16, e0255752 (2021).

76. A. Jacobs, et al., Temperature-dependent gene regulatory divergence underlies local adaptation with gene flow in the Atlantic silverside. Evolution 78, 1133–1149 (2024).

77. A. M. Bolger, M. Lohse, B. Usadel, Trimmomatic: a flexible trimmer for Illumina sequence data. Bioinformatics 30, 2114–2120 (2014).

78. B. Langmead, S. L. Salzberg, Fast gapped-read alignment with Bowtie 2. Nat. Methods. 9, 357–359 (2012).

79. H. Li, et al., The sequence alignment/map format and SAMtools. Bioinformatics 25, 2078–2079 (2009).

80. A. McKenna, et al., The genome analysis toolkit: A MapReduce framework for analyzing next-generation DNA sequencing data. Genome Res. 20, 1297–1303 (2010).

81. T. S. Korneliussen, A. Albrechtsen, R. Nielsen, ANGSD: Analysis of next generation sequencing data. BMC Bioinform. 15, 356 (2014).

82. J. Meisner, A. Albrechtsen, Inferring population structure and admixture proportions in low-depth NGS Data. Genetics 210, 719–731 (2018).

83. A. R. Quinlan, I. M. Hall, BEDTools: a flexible suite of utilities for comparing genomic features. Bioinformatics 26, 841–842 (2010).

84. G. Bhatia, N. Patterson, S. Sankararaman, A. L. Price, Estimating and interpreting F_ST:_ The impact of rare variants. Genome Res. 23, 1514–1521 (2013).

85. D. A. Marques, F. C. Jones, F. D. Palma, D. M. Kingsley, T. E. Reimchen, Experimental evidence for rapid genomic adaptation to a new niche in an adaptive radiation. Nat. Ecol. Evol. 2, 1128–1138 (2018).

86. A. Auton, et al., Genetic recombination is targeted towards gene promoter regions in dogs. PLoS Genet. 9, e1003984 (2013).

87. S. Singhal, et al., Stable recombination hotspots in birds. Science 350, 928–932 (2015).

88. A. F. Shanfelter, S. L. Archambeault, M. A. White, Divergent fine-scale recombination landscapes between a freshwater and marine population of threespine stickleback fish. Genome Biol. Evol. 11, 1573–1585 (2019).

89. R. C. Team, R: 2019. A Language and Environment for Statistical Computing version 3 (2020).

90. C. B. Krimbas, J. R. Powell, Drosophila inversion polymorphism (CRC press, 1992).

91. T. D. Petes, Meiotic recombination hot spots and cold spots. Nat. Rev. Genet. 2, 360–369 (2001).

92. Y. Mansour, A. Chateau, A.-S. Fiston-Lavier, BREC: an R package/Shiny app for automatically identifying heterochromatin boundaries and estimating local recombination rates along chromosomes. Bmc Bioinformatics 22, 396 (2021).

93. G. Benson, Tandem repeats finder: a program to analyze DNA sequences. Nucleic Acids Res. 27, 573–580 (1999).

94. J. Meyne, R. L. Ratliff, R. K. Moyzis, Conservation of the human telomere sequence (TTAGGG)n among vertebrates. Proc. Natl. Acad. Sci. 86, 7049–7053 (1989).

